# Long range channelrhodopsin-assisted circuit mapping of inferior colliculus neurons with blue and red-shifted channelrhodopsins

**DOI:** 10.1101/756957

**Authors:** David Goyer, Michael T. Roberts

**Author notes:** Corresponding author: Michael T. Roberts. Email addresses of co-author: David Goyer.

## Abstract

When investigating neural circuits, a standard limitation of the in vitro patch clamp approach is that axons from multiple sources are often intermixed, making it difficult to isolate inputs from individual sources with electrical stimulation. However, by using channelrhodopsin assisted circuit mapping (CRACM) this limitation can now be overcome. Here, we report a method to use CRACM to map ascending inputs from lower auditory brainstem nuclei and commissural inputs to an identified class of neurons in the inferior colliculus (IC), the midbrain nucleus of the auditory system. In the IC, local, commissural, ascending, and descending axons are heavily intertwined and therefore indistinguishable with electrical stimulation. By injecting a viral construct to drive expression of a channelrhodopsin in a presynaptic nucleus, followed by patch clamp recording to characterize the presence and physiology of channelrhodopsin-expressing synaptic inputs, projections from a specific source to a specific population of IC neurons can be mapped with cell type-specific accuracy. We show that this approach works with both Chronos, a blue light-activated channelrhodopsin, and ChrimsonR, a red-shifted channelrhodopsin. In contrast to previous reports from the forebrain, we find that ChrimsonR is robustly trafficked down the axons of dorsal cochlear nucleus principal neurons, indicating that ChrimsonR may be a useful tool for CRACM experiments in the brainstem. The protocol presented here includes detailed descriptions of the intracranial virus injection surgery, including stereotaxic coordinates for targeting injections to the dorsal cochlear nucleus and IC of mice, and how to combine whole cell patch clamp recording with channelrhodopsin activation to investigate long-range projections to IC neurons. Although this protocol is tailored to characterizing auditory inputs to the IC, it can be easily adapted to investigate other long-range projections in the auditory brainstem and beyond.

**SUMMARY:** Channelrhodopsin-assisted circuit mapping (CRACM) is a precision technique for functional mapping of long-range neuronal projections between anatomically and/or genetically identified groups of neurons. Here, we describe how to utilize CRACM to map auditory brainstem connections, including the use of a red-shifted opsin, ChrimsonR.

## INTRODUCTION

Synaptic connections are critical to neural circuit function, but the precise topology and physiology of synapses within neural circuits is often difficult to probe experimentally. This is because electrical stimulation, the traditional tool of cellular electrophysiology, indiscriminately activates axons near the stimulation site, and in most brain regions, axons from different sources (local, ascending, and/or descending) intertwine. However, by using channelrhodopsin assisted circuit mapping (CRACM)^1,2^, this limitation can now be overcome^3^. Channelrhodopsin (ChR2) is a light activated, cation-selective ion channel originally found in the green alga Chlamydomonas reinhardtii. ChR2 can be activated by blue light of a wavelength around 450-490 nm, depolarizing the cell through cation influx. ChR2 was first described and expressed in Xenopus oocytes by Nagel and colleagues^4^. Shortly after that, Boyden and colleagues^5^ expressed ChR2 in mammalian neurons and showed that they could use light pulses to reliably control spiking on a millisecond timescale, inducing action potentials ∼10 ms after activation of ChR2 with blue light. Optogenetic channels with even faster kinetics have been found recently, e.g. Chronos^6^.

The basic approach to a CRACM experiment is to transfect a population of putative presynaptic neurons with a recombinant adeno-associated virus (rAAV)-vector that carries the genetic information for a channelrhodopsin. Transfection of neurons with rAAV leads to the expression of the encoded channelrhodopsin. Typically, the channelrhodopsin is tagged with a fluorescent protein like GFP (Green Fluorescent Protein) or tdTomato (red fluorescence), so that transfection of neurons in the target region can easily be confirmed with fluorescence imaging. Because rAAVs are non-pathogenic, have a low inflammatory potential and long lasting gene expression^7,8^, they have become a standard technique to deliver channelrhodopsins to neurons. If, after transfection of a putative presynaptic population of neurons, activation of a channelrhodopsin through light flashes elicits postsynaptic potentials or currents in the target neurons, this is evidence of an axonal connection from the transfected nucleus to the recorded cell. Because severed axons in brain slice experiments can be driven to release neurotransmitter through channelrhodopsin activation, nuclei that lie outside of the acute slice but send axons into the postsynaptic brain region (the IC in our case) can be identified with CRACM. The power of this technique is that the connectivity and physiology of identified long range synaptic inputs can be directly investigated.

In addition to channelrhodopsins that are excitable by blue light, investigators have recently identified several red-shifted channelrhodopsins^9,10^, including Chrimson and its faster analog ChrimsonR, both of which are excited with red light of ∼660 nm^6^. Red-shifted opsins are of interest because red light penetrates tissue better than blue light, and red light may have a lower cytotoxicity than blue light^10–12^. Red-shifted channelrhodopsins also open up the possibility of dual color CRACM experiments, where the convergence of axons from different nuclei on the same neuron can be tested in one experiment^6,13,14^. However, current red-shifted opsins often exhibit unwanted cross-activation with blue light^15–17^, making two color experiments difficult. In addition, some reports have indicated that ChrimsonR undergoes limited axonal trafficking, which can make it challenging to use ChrimsonR for CRACM experiments^16,17^.

Nearly all ascending projections from the lower auditory brainstem nuclei converge in the inferior colliculus (IC), the midbrain hub of the central auditory pathway. This includes projections from the cochlear nucleus (CN)^18,19^, most of the superior olivary complex (SOC)^20^, and the dorsal (DNLL) and ventral (VNLL) nuclei of the lateral lemniscus^21^. Additionally, a large descending projection from the auditory cortex terminates in the IC^18–22^, and IC neurons themselves synapse broadly within the local and contralateral lobes of the IC^23^. The intermingling of axons from many sources has made it difficult to probe IC circuits using electrical stimulation^24^. As a result, even though neurons in the IC perform computations important for sound localization and the identification of speech and other communication sounds^25,26^, the organization of neural circuits in the IC is largely unknown. We recently identified VIP neurons as the first molecularly-identifiable neuron class in the IC^27^. VIP neurons are glutamatergic stellate neurons that project to several long range targets, including the auditory thalamus and superior colliculus. We are now in a position to determine the sources and function of local and long range inputs to VIP neurons and to determine how these circuit connections contribute to sound processing.

The protocol presented here is tailored to investigating synaptic inputs to VIP neurons in the IC of mice, specifically from the contralateral IC and the DCN (Fig. 1). The protocol can be easily adapted to different sources of input, a different neuron type or a different brain region altogether. We also show that ChrimsonR is an effective red-shifted channelrhodopsin for long range circuit mapping in the auditory brainstem. However, we demonstrate that ChrimsonR is strongly activated by blue light, even at low intensities, and thus, to combine ChrimsonR with Chronos in two-color CRACM experiments, careful controls must be used to prevent cross activation of ChrimsonR.

**Figure 1:**
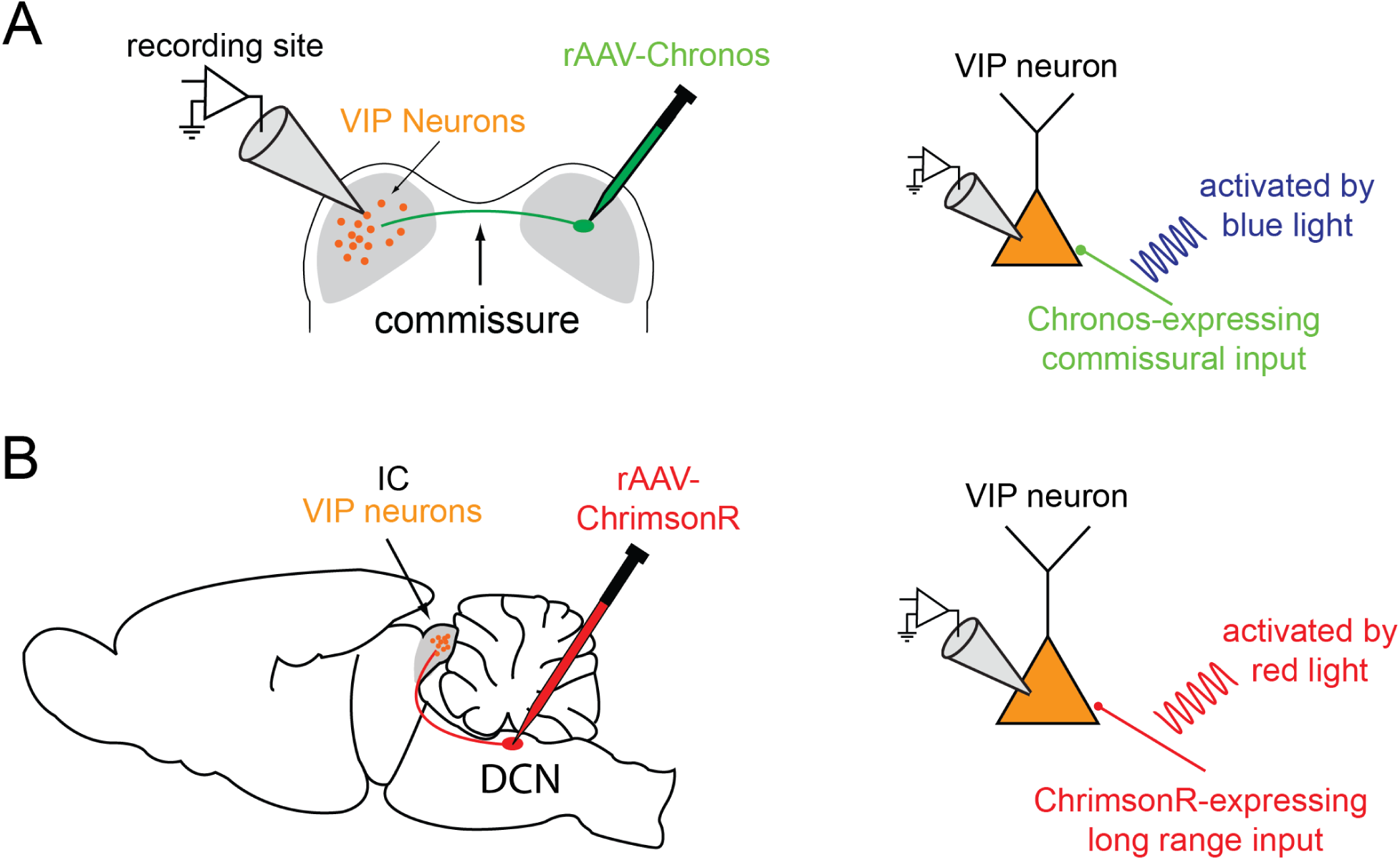
rAAV injection sites and experimental setup. **(A)** Injection site and experimental setup to investigate commissural projections. Left: An rAAV construct (e.g. rAAV1.Syn.Chronos-GFP.WPRE.bGH) is injected into the right IC and targeted recordings are performed in the contralateral IC. Right: During patch clamp recordings, Chronos-transfected fibers are excited with blue light to elicit postsynaptic potentials. **(B)** Injection site and experimental setup to investigate long-range projections from DCN to IC. Left: The ChrimsonR construct is injected into the right DCN and targeted recordings are performed in the contralateral IC. Right: During patch clamp recordings, ChrimsonR-transfected fibers are excited with red light to record ChrimsonR evoked postsynaptic potentials.

## PROTOCOL

### 1. Surgery preparations

1.1. Surgeries need to be done in aseptic conditions. Autoclave / sterilize all surgery tools and materials before surgery, sanitize the surgery area (spray and wipe down with 70% ethanol), and place sterile towel drapes to cover the surgery area. Wear surgery gown and mask for the surgery.
1.2. Ready recovery cage. Remove cage bedding to limit risk of asphyxiation. Put heating pad under cage. Provide a food and water source (e.g. DietGel).
1.3. Pull glass capillary for nano injector on a pipette puller. Cut or break off tip to obtain an opening approximately 5 µm in diameter. Bevel capillary tip to an approximately 30° angle to improve tissue penetration and reduce clogging. Backfill capillary with mineral oil and insert into Drummond Nanoject (or other nanoliter injector).
1.4. Obtain an aliquot of the desired channelrhosopdin-encoding rAAV and dilute to desired titer using sterile PBS. We have found that serotype 1 rAAVs work well for transfection of auditory brainstem nuclei. Specifically, rAAV1.Syn.Chronos-GFP.WPRE.bGH (blue light-activated channelrhodopsin) and rAAV1.Syn.ChrimsonR-tdTomato.WPRE.bGH (red-shifted channelrhodopsin) consistently yield the high expression levels and good long-range axonal trafficking of channelrhodopsins needed for CRACM experiments.
1.5. Follow injector instructions to front fill capillary with 1 – 3 µl of rAAV in sterile PBS.

### 2. Surgery

2.1. Put animal into induction chamber and induce anesthesia with 3% isoflurane in oxygen delivered via a calibrated isoflurane vaporizer. Observe mouse until breathing becomes deep and slow and toe pinch reflex is absent, about 3 – 5 minutes.
2.2. Transfer animal to stereotaxic frame. Secure animal’s head by putting its mouth on a palate bar with a gas anesthesia mask and by positioning non-perforating ear bars in both ear canals.
2.3. Insert rectal temperature probe and switch on homeostatic temperature controller.
2.4. Apply ophthalmic ointment to prevent eyes from drying out.
2.5. Administer preemptive analgesic (e.g. subcutaneous injection of 5 mg/kg carprofen).
2.6. Adjust isoflurane to 1 - 2.5%, according to depth of anesthetized state. Monitor temperature, breathing and color of mucous membranes at least every 15 minutes during the procedure.
2.7. Shave scalp with electric clippers. Disinfect scalp with three alternating swabs of povidone-iodine and 70% ethanol.
2.8. Make an incision in the scalp along the midline starting between the ears and continuing rostral to the eyes, exposing the lambda and bregma sutures. Push skin to the side and remove periosteum from exposed bone if necessary.
2.9. Mark the lambda suture with a surgical marker, position the tip of the nanoinjector so that it is just touching lambda, and zero the micromanipulator coordinates. Use the nanoinjector tip and micromanipulator to measure the difference in elevation between the lambda and bregma sutures. Adjust palate bar height to bring lambda and bregma to within ±100 µm height difference.
2.10. Map the injection site using the nanoinjector tip and micromanipulator coordinate system and mark the site with a surgical marker. To inject the IC or DCN of P21 – P30 animals, use coordinates relative to the lambda suture, as shown in Table 1:
2.11. Use a micromotor drill with a 0.5 mm drill burr to perform a craniotomy over the injection site.
2.12. To ensure broad transfection of neurons in the target nucleus, injections are made at various depths into the tissue (Table 1, Z coordinates), and, in the case of larger brain regions like the IC, injections are made over the course of two or more penetrations at different X and Y coordinates (Table 1, Right IC penetration 1 and Right IC penetration 2).
2.13. Perform injections. For IC injections, deposit 20 nl of virus in intervals of 250 µm along the Z axis (injection depth) between 2250 µm and 1750 µm depth. For DCN injections, deposit 20 nl virus at a depth of 4750 µm and 4550 µm, respectively. After injection at each Z coordinate, wait 2 – 3 minutes before moving injector to next Z coordinate. This will allow time for virus to diffuse away from injection site, reducing the probability that virus will be sucked up the injection tract when the nanoinjector is repositioned. After last injection in a penetration, wait 3 – 5 minutes before retracting nanoinjector from brain.
2.14. When the nanoinjector is removed from the brain between penetrations and between animals, eject a small volume of virus from the tip to check that tip has not clogged.
2.15. After injections, use sterile PBS to wet the cut edges of the scalp and then gently move the skin back towards the midline. Close the wound with simple interrupted sutures using 6-0 (0.7 metric) nylon sutures. Apply 0.5 – 1 ml 2% lidocaine jelly to the wound.
2.16. Remove ear bars and temperature probe, turn off isoflurane, remove mouse from the palate bar and transfer it to the recovery cage. Monitor recovery closely. Once the animal is fully awake, moving around, and showing no signs of pain or distress, transfer it back into its cage and return the cage to the vivarium.
2.17. If surgeries will be performed on multiple animals in one day, use a hot bead sterilizer to sanitize surgery tools before next surgery.

**Table 1:** Stereotaxic coordinates for rAAV injection into IC and DCN. Coordinates are relative to lambda in µm.

|  | X (lateral) | Y (caudal) | Z (depth) |
| --- | --- | --- | --- |
| Right IC penetration 1 | 1000 $\mu\text{m}$ | -900 $\mu\text{m}$ | 2250-1500 $\mu\text{m}$ (250 $\mu\text{m}$ increments) |
| Right IC penetration 2 | 1250 $\mu\text{m}$ | -900 $\mu\text{m}$ | 2250-1750 $\mu\text{m}$ (250 $\mu\text{m}$ increments) |
| Right DCN | 2155 $\mu\text{m}$ | -1325 $\mu\text{m}$ | 4750 $\mu\text{m}$, 4550 $\mu\text{m}$ |

### 3. Surgery follow up

3.1. Check animals daily for wound closure, infection, or signs of pain or distress over the next 10 days, adhering to the institution’s animal care guidelines.
3.2. Wait 3 – 4 weeks before using animals in experiments to allow optimal expression of the channelrhodopsins.

### 4. Brain slice preparation and confirmation of injection target

4.1. For CRACM, acutely prepared brain slices from transfected animals are used in standard in vitro electrophysiology experiments, described here only briefly (see Goyer et al., 2019 for a more detailed description).
4.2. Prepare artificial cerebrospinal fluid (ACSF) containing (in mM): 125 NaCl, 12.5 D-glucose, 25 NaHCO_3_, 3 KCl, 1.25 NaH_2_PO_4_, 1.5 CaCl_2_, 1 MgSO_4_. Bubble ACSF to a pH of 7.4 with 5% CO_2_ in 95% O_2_.
4.3. Perform all following steps, including in vitro electrophysiology, in near-darkness or red light to limit activation of channelrhodopsins.
4.4. Deeply anesthetize mouse with isoflurane and decapitate it quickly. Dissect the brain quickly in ∼34 °C ACSF.
4.5. Cut coronal slices (200 – 250 µm) containing the IC in ∼34 °C ACSF with a vibrating microtome and incubate the slices at 34 °C for 30 minutes in a holding chamber filled with ACSF bubbled with 5% CO_2_ in 95% O_2_. After incubation, store slices at room temperature until used for recordings.
4.6. If the injection target was not the IC, cut additional coronal slices of the injected brain region and check the transfection of the target nucleus under a fluorescence microscope. If there is no transfection in the target nucleus or additional transfection in different brain regions, do not continue with experiment.

### 5. In vitro recording & CRACM experiment

5.1. Pull electrodes from borosilicate glass to a resistance of 3.5 – 4.5 MΩ. The electrode internal solution should contain (in mM): 115 K-gluconate, 7.73 KCl, 0.5 EGTA, 10 HEPES, 10 Na_2_ phosphocreatine, 4 MgATP, 0.3 NaGTP, supplemented with 0.1% biocytin (w/v), pH adjusted to 7.3 with KOH and osmolality to 290 mOsm/kg with sucrose.
5.2. To make recordings, use standard patch clamp methods. Place slice in a recording chamber under a fixed stage upright microscope and continuously perfuse with ACSF at ∼2 ml/min. Conduct recordings near physiological temperature (∼34 - 36 °C). Patch neurons under visual control using a suitable patch clamp amplifier. Correct for series resistance, pipette capacitance and liquid junction potential.
5.3. During whole cell recordings, activate Chronos by delivering brief pulses (1 – 5 ms) of 470 nm light or ChrimsonR by brief pulses of 580 nm light. Determine threshold of opsin activation and use a minimal stimulation protocol to elicit postsynaptic potentials. In general, use the shortest stimulus duration that elicits a PSP, and set the optical power to 120% of the threshold power required to elicit PSPs.
5.4. To confirm that the recorded changes in membrane potential are indeed synaptic inputs to the neuron, standard antagonists for excitatory / inhibitory postsynaptic receptors can be washed in during the experiment. To investigate different receptor contributions to a PSP (e.g. NMDA vs AMPA receptors), suitable receptor antagonists can be washed in. For each receptor antagonist, drug effects should reverse after washout.
5.5. The latency, jitter, and reliability of PSPs should be used to confirm that light-activated synaptic inputs originate from direct, optical activation of synapses on the recorded neuron, as opposed to activation of channelrhodopsin-expressing synapses on an intervening neuron that synapses on the recorded neuron. In general, low latency (< 2 ms), low jitter (<1 ms standard deviation in latency), and high reliability (>50%) indicate a direct synaptic connection from the channelrhodopsin expressing presynaptic neuron to the recorded neuron.

## REPRESENTATIVE RESULTS

Stereotaxic injection of AAV1.Syn.Chronos-GFP.WPRE.bGH into the right IC using coordinates shown in Table 1 results in strong Chronos-GFP expression in the right IC (Figure 2A). Visual inspection of Chronos-GFP fluorescence indicates that most of the somata labeled in the right IC are located in the central nucleus of the IC (ICc), but labeled somata are sometimes also present in the dorsal cortex of the IC (ICd) and occasionally in the lateral cortex of the IC (IClc). The targeting and extent of transfection should be checked for every animal used in an experiment, as expression of channelrhodopsins in non-targeted regions can lead to false positives. To achieve broader or more restricted expression of Chronos, the amount of deposited virus as well as the stereotaxic coordinates can be easily adjusted to achieve the desired outcome.

**Figure 2:**
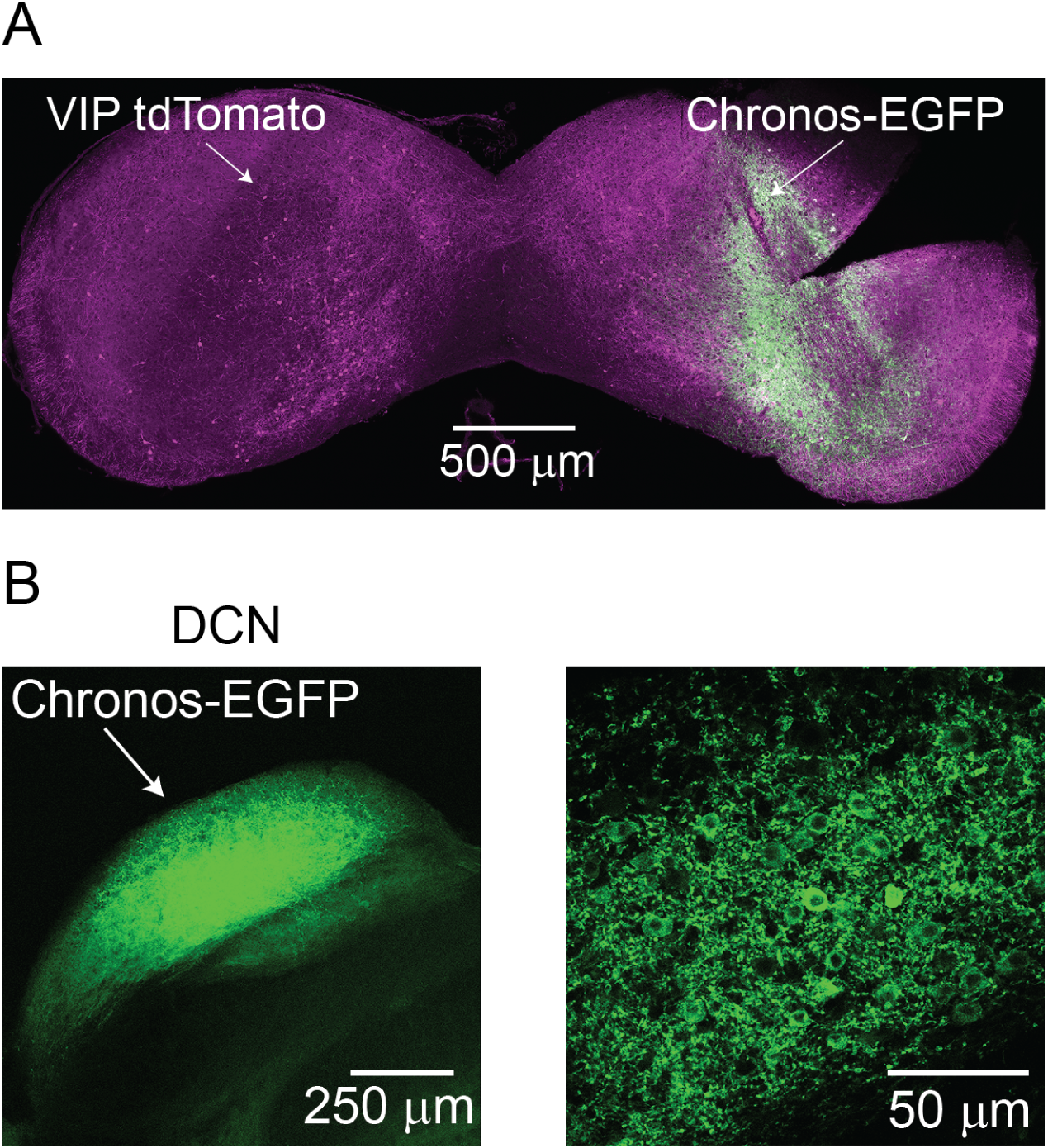
Chronos-GFP expression after injections into IC and DCN. **(A)** Image of coronal IC slice showing tdTomato-labeled VIP neurons throughout the IC and Chronos-GFP expression in somata localized to the injection sites in the right IC. **(B)** Image of coronal brainstem slice showing Chronos-GFP expression in the DCN after rAAV-Chronos-GFP injection. Left: Low magnification image of the DCN, showing transfected DCN and GFP-positive fibers entering the dorsal acoustic stria. Right: High magnification image showing Chronos-transfected cell bodies in the DCN.

Stereotaxic injection of AAV1.Syn.Chronos-GFP.WPRE.bGH in the DCN (see coordinates in Table 1) results in strong transfection of DCN neurons (Figure 2B). To confirm selective transfection of the DCN, the brainstem of every animal should be sliced to verify that GFP expression was present and limited to the DCN. If there is no transfection or if there is considerable expression of GFP in the auditory nerve or VCN, recordings should not be performed. Using the coordinates shown in Table 1 with a total injection volume of 40 nl, Chronos-GFP expression will be limited to the DCN in most cases. GFP-labeled axons are present in the left (contralateral) ICc 3 weeks after injection for both IC and DCN injection sites.

To test the long range trafficking of ChrimsonR, AAV1.Syn.ChrimsonR-tdTomato.WPRE.bGH was injected into the right DCN, using the same coordinates as with Chronos injections. ChrimsonR injection led to strong expression in the DCN, with tdTomato fluorescence visible in cells and fibers (Figure 3A). In the contralateral ICc, fibers strongly labeled with tdTomato were clearly visible after 3 weeks, demonstrating the long-range trafficking capability of the ChrimsonR-tdTomato construct when injected into auditory brainstem nuclei (Figure 3B). Optical activation of ChrimsonR elicited EPSPs in IC VIP neurons (Figure 5B), indicating that ChrimsonR is a useful tool for long-range CRACM experiments when the experimental parameters demand the use of red light instead of blue light. However, we found that ChrimsonR was readily activated with blue light, showing the same threshold for blue light activation as Chronos (Figure 5). The sensitivity of ChrimsonR to blue light means that special care must be taken to distinguish between inputs transfected with ChrimsonR or Chronos in the same animal^13^.

**Figure 3:**
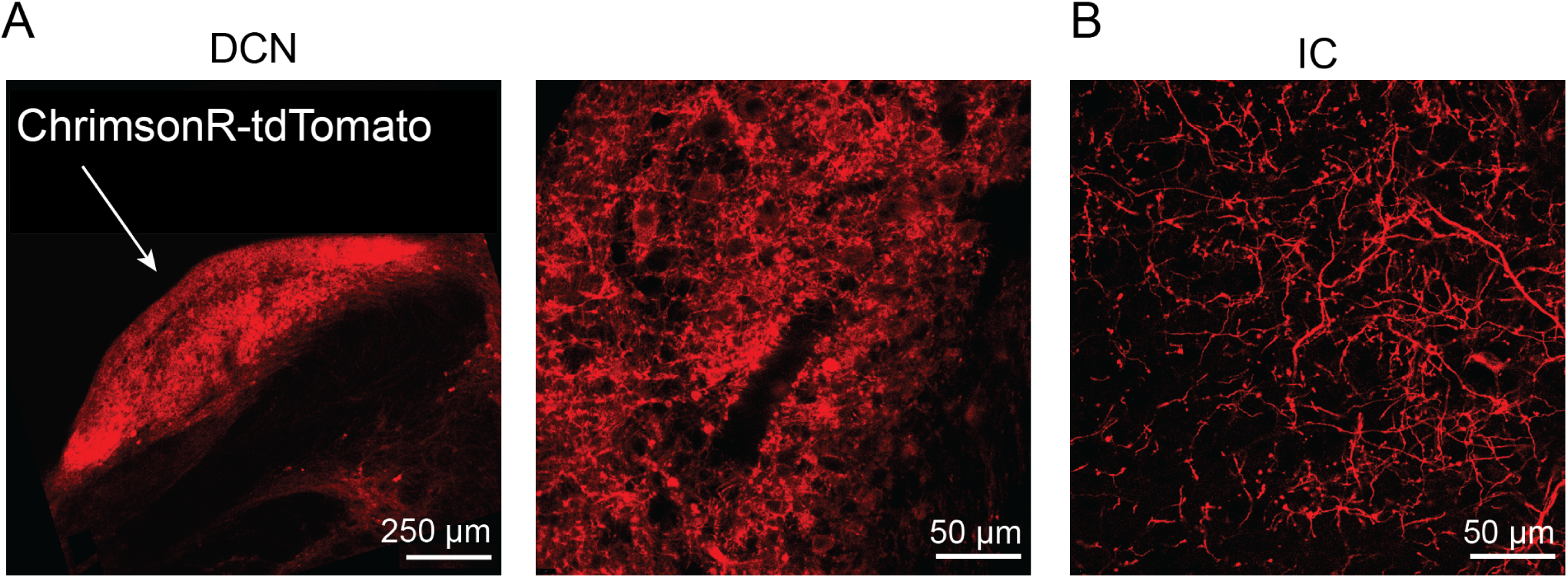
ChrimsonR-tdTomato expression in the DCN and DCN projections to the IC. **(A)** Image of a coronal brainstem slice from a mouse in which the right DCN was injected with rAAV1.Syn.ChrimsonR-tdTomato.WPRE.bGH. Left: Low magnification image of ChrimsonR expression in the right DCN. Middle: High magnification image of right DCN showing strong transfection of neurons with ChrimsonR. **(B)** High magnification image of ChrimsonR-transfected DCN axons in the IC.

**Figure 4:**
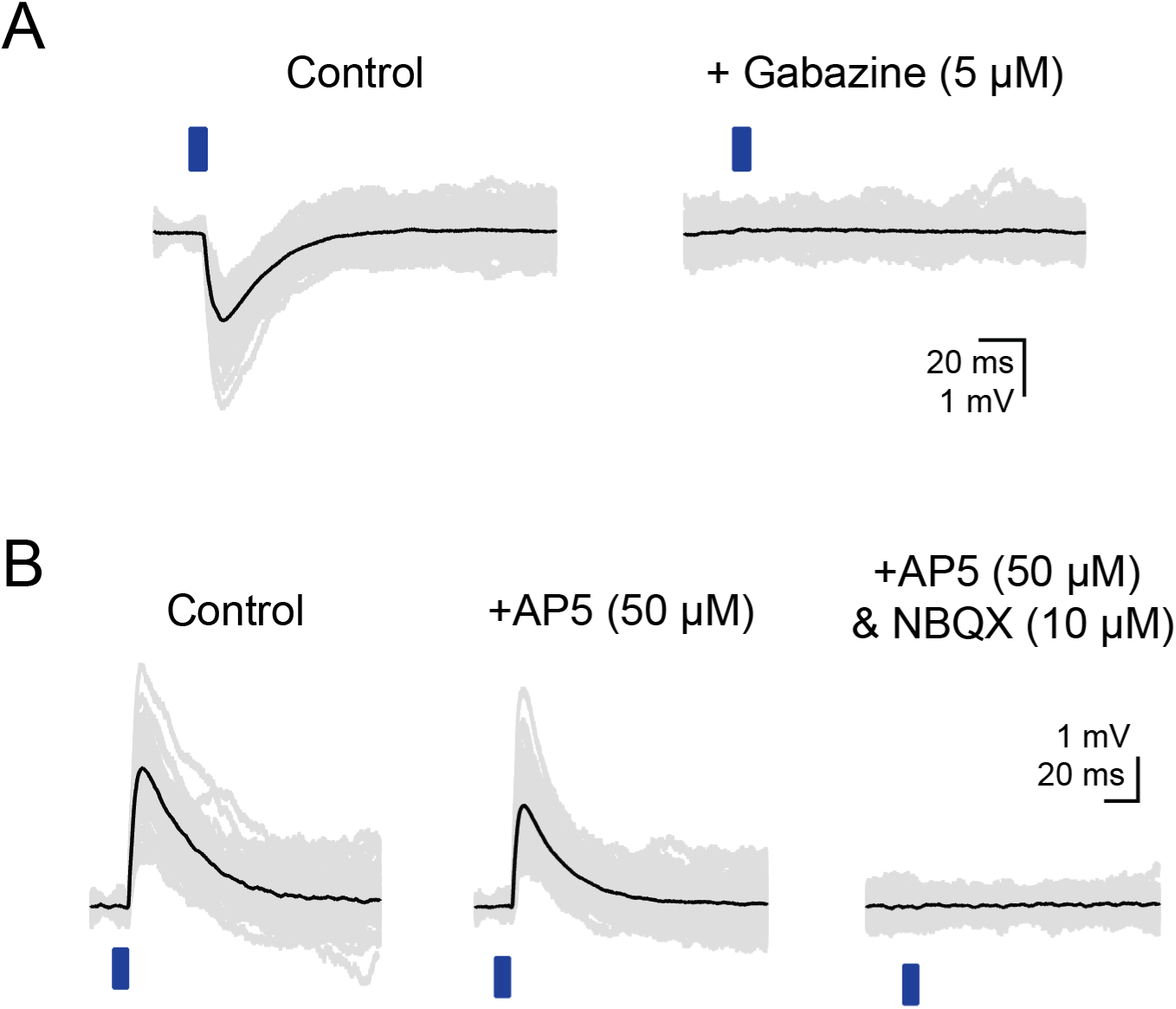
Characterization of light-evoked PSPs recorded in VIP neurons. **(A)** Optogenetically-evoked IPSPs recorded from VIP neurons in the ICc contralateral to the AAV injection site. IPSPs were evoked by 2 – 5 ms blue light flashes (left) while EPSPs were blocked by 10 µm NBQX and 50 µm D-AP5. IPSPs were abolished by gabazine (right). **(B)** Optogenetically-evoked EPSPs recorded from VIP neurons in the ICc contralateral to the AAV injection site. EPSPs were evoked by 2 – 5 ms blue light flashes (left) while IPSPs were blocked with 1 µm strychnine and 5 µm gabazine. Wash-in of 50 µm D-AP5 significantly reduced the halfwidth and decay time constant of light-evoked EPSPs (middle). Wash-in of 10 µm NBQX abolished the remaining EPSP (right). From Goyer et al., 2019.

**Figure 5:**
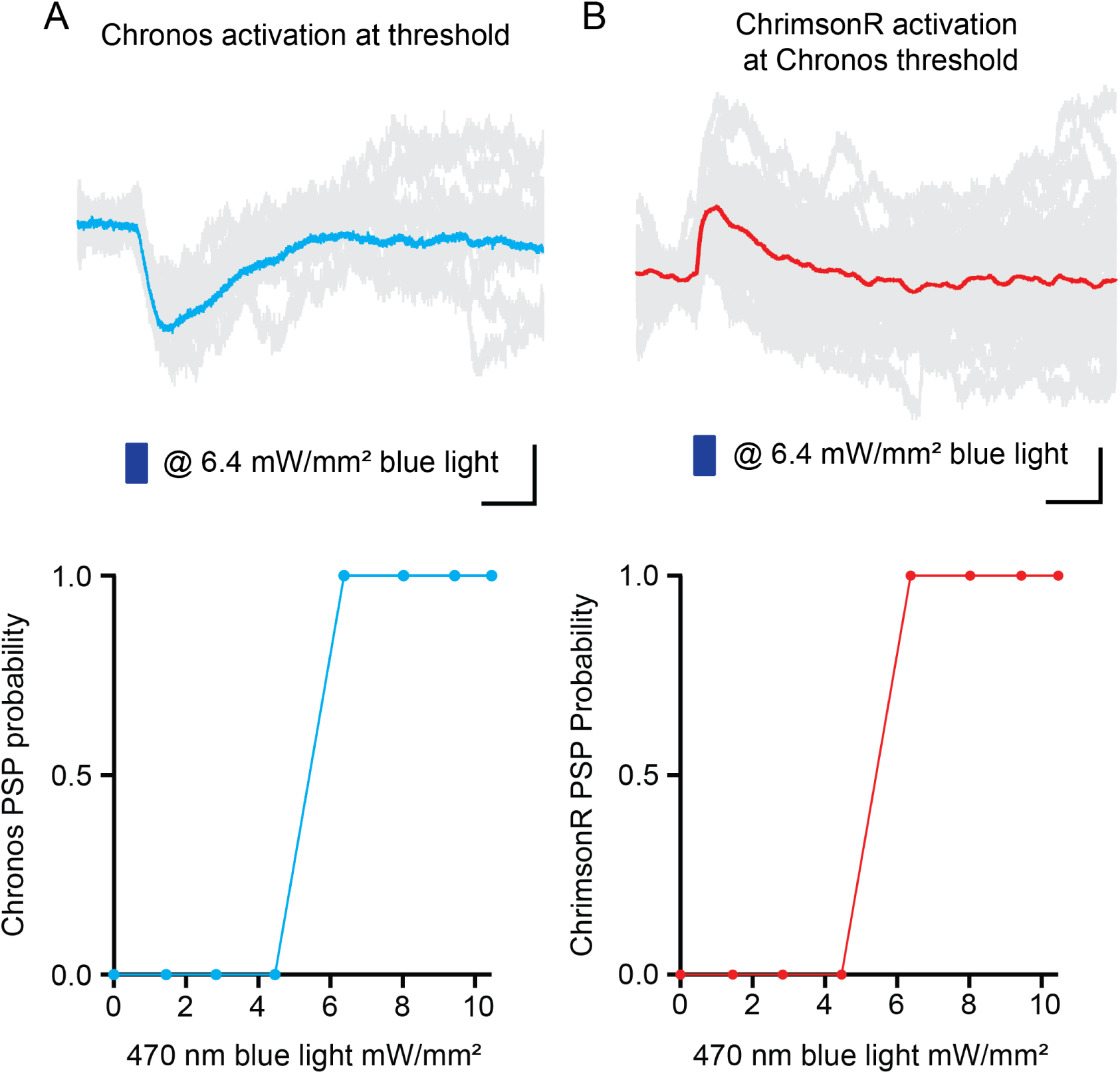
Activation of ChrimsonR by low levels of blue light. **(A**) Activation of Chronos in commissural projections by pulses of 470 nm blue light. Top: Original traces (light grey) and average (cyan) of Chronos-driven IPSPs, evoked at an optical power slightly above the threshold for Chronos activation (scale bar shows 20 ms / 0.5 mV). Bottom: Relationship between optical power at 470 nm and the probability of observing a Chronos-evoked PSP. **(B)** Activation of ChrimsonR with 470 nm blue light. Top: Original traces (light grey) and average (red) of ChrimsonR-driven EPSPs, evoked with 470 nm blue light at the same optical power used in **A, top** (scale bar shows 20 ms / 0.5 mV). Bottom: Relationship between optical power at 470 nm and the probability of observing a ChrimsonR-driven PSP. Note that the threshold for blue light activation of ChrimsonR PSPs was identical to the threshold for eliciting Chronos PSPs.

When targeting recordings to VIP neurons in the contralateral (left) ICc after injections into the right IC, blue light flashes elicited excitatory postsynaptic potentials (EPSPs) or inhibitory postsynaptic potentials (IPSPs). This confirms commissural projections to VIP neurons. To analyze commissural EPSPs and IPSPs separately, we used receptor antagonists to block IPSPs during EPSP recordings, and vice versa. Representative EPSPs and IPSPs recorded during CRACM experiments are shown in Figure 4. IPSPs were observed in 6 out of 12 tested ICc VIP neurons. IPSPs were small (1.53 mV ± 0.96 mV) and had moderate 10 – 90% rise times (7.8 ms ± 2.1 ms), halfwidths (15.1 ms ± 6.8 ms) and decay time constants (32.4 ms ± 17.0 ms) (Figure 4A, left). IPSPs were completely abolished by the GABA_A_ receptor antagonist gabazine (5 µM, Figure 4A, right; n = 6). EPSPs were observed in 11 out of 27 ICc VIP neurons tested. EPSPs were also small (1.52 mV ± 1.08 mV) and had moderate 10 – 90% rise times (8.3 ms ± 4.3 ms), halfwidths (19.6 ms ± 7.6 ms) and decay time constants (43.5 ms ± 16.8 ms) (Figure 4B “Control”). Adding the NMDA receptor antagonist D-AP5 to the bath significantly reduced the halfwidth of EPSPs (14.3 ms ± 4.7 ms, p = 0.006) and revealed a trend toward reducing the rise time (6.3 ms ± 1.6 ms, p = 0.09) and decay time constant (30.6 ms ± 7.3 ms, p = 0.06) of EPSPs (ANOVA for repeated measurements with Tukey post-hoc test). The remainder of the EPSP was completely blocked by the AMPA receptor antagonist NBQX (Figure 4B “+ AP5 & NBQX”).

Recordings of VIP neurons in the left ICc after DCN injections revealed EPSPs evoked by blue light flashes, confirming synaptic inputs from the DCN to VIP neurons in the IC. DCN CRACM experiments were conducted with GABAergic and glycinergic blockers in the bath to block spontaneous IPSPs. We found that 2 – 5 ms pulses of blue light elicited EPSPs in 19 of 25 neurons tested. Light-evoked EPSPs had moderate amplitudes (2.85 mV ± 2.98 mV) and relatively slow rise times (4.2 ms ± 1.3 ms), halfwidths (20.6 ms ± 14.4 ms) and decay time constants (22.0 ms ± 6.7 ms) (n = 6 cells, data not shown).

## DISCUSSION

We have found that CRACM is a powerful technique for identifying and characterizing long range synaptic inputs to neurons in the mouse IC. Following the protocol detailed here, we achieved robust transfection of neurons in the DCN and IC as well as reliable axonal trafficking of Chronos and ChrimsonR to synaptic terminals in the IC. Additionally, we demonstrated that this technique enables the measurement and analysis of postsynaptic events, including PSP amplitude, halfwidth, decay time, and receptor pharmacology. Our experience suggests that this approach can be readily adapted to perform functional circuit mapping experiments throughout the auditory brainstem and beyond.

Overall, the specificity of optogenetic circuit mapping provides a distinct advantage over electrical stimulation of axons. Viral transfections provide the ability to spatially and molecularly restrict the expression of channelrhodopsins to a targeted population of presynaptic neurons. In contrast, electrical stimulation cannot differentiate between axons originating from different presynaptic nuclei when those axons are intermingled, as is the case in most brain regions. Electrical stimulation can also initiate both orthodromic and antidromic spikes, further complicating matters when a distal stimulation site contains axons originating from neurons located near the recording site.

To ensure stable expression and good axonal trafficking of a channelrhodopsin, choosing the right viral vector and serotype is paramount. We found that stable expression of Chronos in IC neurons was achieved with a serotype 1 rAAV including a Chronos or ChrimsonR construct combined with a woodchuck hepatitis posttranscriptional regulatory element (WPRE) and bovine growth hormone (BGH) polyadenylation signal. rAAV serotype 5 failed to produce functional opsins in the IC, but was functional in the DCN (data not shown). The rigorous validation of injection coordinates for every experiment and ensuring that opsin expression is restricted to the targeted brain region is similarly important. A meaningful identification of inputs is only possible if expression of the opsin is carefully checked and documented for every experiment.

The AAV1.ChrimsonR construct used in the above protocol yielded stable expression and good axonal trafficking of the protein, even in long range projections from the DCN to the IC (Fig. 3). This makes ChrimsonR a suitable opsin for (long range) circuit mapping. A red-shifted opsin like ChrimsonR can be useful if the experimental parameters require light penetration deep into the tissue, but the experimenter must be aware that all currently-available red-shifted opsins show some cross-activation with blue light. Although some studies have argued that ChrimsonR and Chronos may be separately activated^6,14,16^, our data suggest that great care must be taken with this approach. A recent report details additional methods that should be used if attempting to separate red-shifted opsins from Chronos^13^. Therefore, when using ChrimsonR and Chronos in two color CRACM experiments, carefully designed control experiments need to be executed to ensure clear separation of blue and red shifted opsin activation.

## ACKNOWLEDGMENTS

This work was supported by a Deutsche Forschungsgemeinschaft Research Fellowship (GO 3060/1–1, project number 401540516, to DG) and National Institutes of Health grant R56 DC016880 (MTR).

## DISCLOSURES

The authors have nothing to disclose.

